# Hough transform implementation to evaluate the morphological variability of the moon jellyfish (*Aurelia* spp.)

**DOI:** 10.1101/2020.03.11.986984

**Authors:** Céline Lacaux, Agnès Desolneux, Justine Gadreaud, Bertrand Martin-Garin, Alain Thiéry

## Abstract

Variations of the animal body plan morphology and morphometry can be used as prognostic tools of their habitat quality. The potential of the moon jellyfish (*Aurelia* spp.) as a new model organism has been poorly tested. However, as a tetramerous symmetry organism, it exhibits some variations in radial symmetry number. A pertinent list of morphological – number of gonads – and morphometric characteristics – e.g. ratio of the gonads area on the umbrella area – has been established to describe the morphology of 19 specimens through an image analysis. The method uses for the first time the Hough transform to approximate the gonads and the umbrella by ellipses and automatically extracts the morphometric data. A statistical comparison has been done to compare the morphometric characteristics of tetramerous jellyfish and of jellyfish with 5 gonads: it is only provided as a first step for testing biological hypotheses, since the small size of the data set leads to relativize its conclusions. It suggests that two parameters are discriminant: distance between the center of the gonads and the center of the umbrella, and the individual variability of the gonad eccentricity, both higher in jellyfish with 5 gonads. Additionally, the relative size of the gonads does not seem to be different between tetramerous and non-tetramerous. Combined to ecotoxicological bioassays to better understand the causes of this developmental alteration, this optimizable method can become a powerful tool in the symmetry description of an *in situ* population.

## Introduction

### Jellyfish proliferations and asymmetric specimens of Aurelia spp

Over the last decades, the proliferation of adult jellyfish has increased worldwide both in intensity and frequency along many marine coastal areas causing harmful societal inconveniences for industry and populations such as reduction of the fishery production, tourism, stinging of swimmers, etc. (Dong *et al*. 2010; Purcell *et al*. 2007; Richardson *et al*. 2009). It is commonly accepted that part of these blooms is a consequence of environmental changes often induced by intensive anthropogenic disturbance (Purcell 2005; Richardson *et al*. 2009) such as eutrophication, overfishing, translocation, habitat modification, etc. (Dong *et al*. 2010; Purcell 2005; Purcell *et al*. 2007; Richardson *et al*. 2009). The moon jellyfish *Aurelia* sp. is a diploblastic Semaeostomeae cnidarian with a worldwide distribution and the most common jellyfish in Europe coastal environments (Yuan *et al*. 2008). Scyphomedusae including moon jellyfish are tetramerous by definition with a stomach divided in four gastro-gonadic pouches by mesenteries in the center of their umbrella (Figure **1**A). However, it can exhibit some variations in radial symmetry number (Brusca and Brusca 2003) (Figure **1**B and C). The proportion of the non-tetramerous specimen in wild population has been estimated around 2% and sparsely varies depending on the location (Gershwin 1999). The public aquariums also notice around 6–15% of non-tetramerous jellyfish in their populations (**Table 1**).

**Table 1.** Rough estimations of non-tetramerous jellyfish proportions in public aquariums – personal observations.

| Public aquariums | Monterey Bay (USA) | Monaco | Barcelona (Spain) | La Rochelle (France) | Vancouver (Canada) |
| --- | --- | --- | --- | --- | --- |
| Dates | 10 August 2013 | 3 November 2014 | 1 June 2016 | 20 December 2016 | 13 August 2017 |
| Non-tetramerous jellyfish proportions | 11,5 %<br>(*n* = 122) | 6 %<br>(*n* = 160) | 15 %<br>(*n* ≈ 80) | 9 %<br>(*n* ≈ 200) | 13 %<br>(*n* = 120) |

In the Berre lagoon at 20 km west of Marseille (France), proliferation events of the moon jellyfish *Aurelia* sp. were observed in brackish environments as those of the summers 2006 and 2008 with a high proportion (≈ 6–7%) of non-tetramerous specimens (Delpy *et al*. 2012). These phenotypic responses may be caused by a disturbance in the developmental process during the strobilation and the morphogenesis of the ephyrae. Indeed, the Berre lagoon is the largest French lagoon of the Mediterranean coast: its environment is affected by chemical pollutions and anthropogenic effluents (Accornero *et al*. 2008; Gadreaud *et al*. 2017; Rigaud *et al*. 2011).

### Symmetry disorders as a biomarker

The body symmetry – *Bauplan* concept *sensu* Brusca and Brusca (2003) – appeared early in the evolution process *ca*. 575 million years ago (Ediacaran age). It can be defined as a balanced distribution of duplicate body part. It is a common characteristic of the eumetazoans; all the animals excepting the sponges, but including the cnidarian phylum. As primitive organisms, each stage of cnidarian life cycle exhibits a vertical polar axis and a tetramerous radial symmetry axis from the center of the oral surface (
Figure 2).

**Figure 1.**
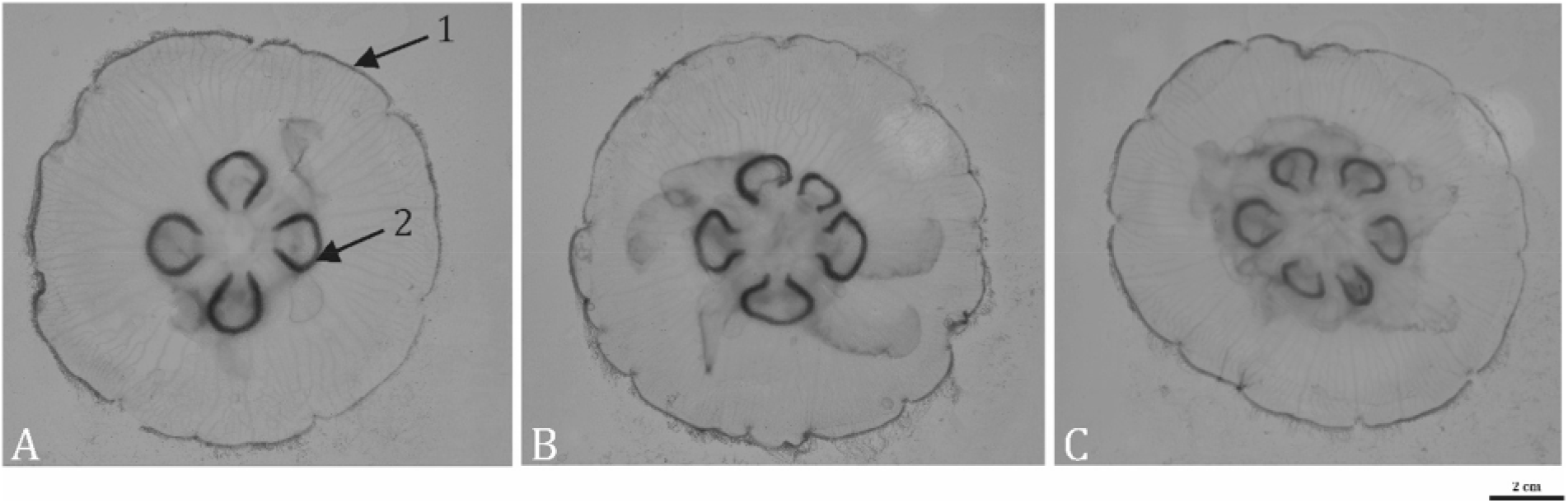
Specimen of *Aurelia* sp. sampled in the Berre lagoon; 1: umbrella; 2: gastro-gonadic pouches (gonads); A) a tetramerous symmetry N = 4 gonads; B) and C) a non-tetramerous symmetry, N = 5, and N = 6 gonads respectively.

**Figure 2.**
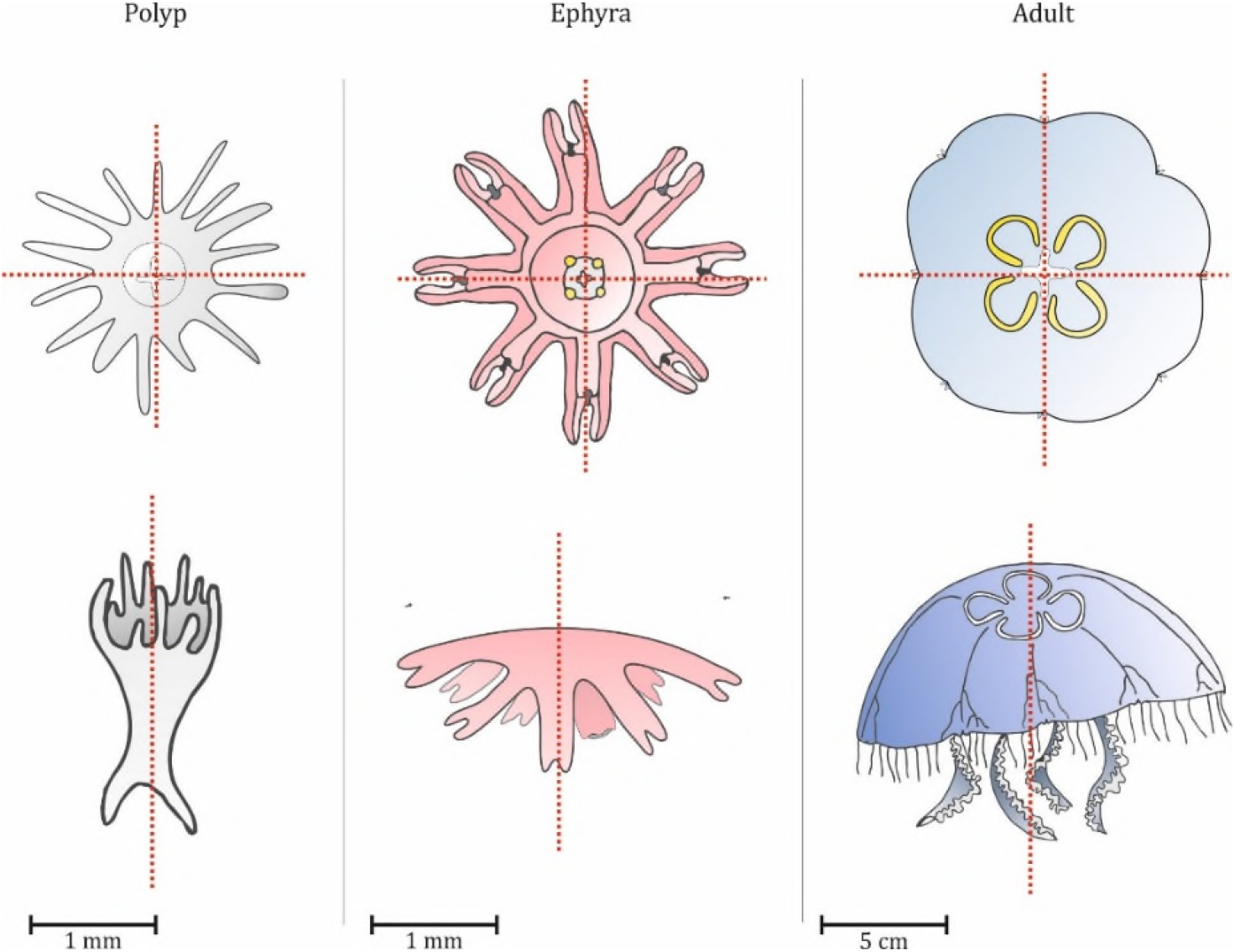
Symmetry at each stage of the *Aurelia* spp. jellyfish: tetramerous radial symmetry (oral face view) and vertical polar axis on polyp, ephyrae and medusae.

The characterization and the quantification of the elliptic characters of the jellyfish (global shape and number of gastro-gonadal pouches of the tetramerous and non-tetramerous specimens) remain a preliminary challenge to provide a database on the proportion of asymmetric jellyfish and to evaluate the evolutionary advantages of reproduction or feeding. During a proliferation event of the moon jellyfish, the specimens can be easily sampled and photographed directly on the boat. Then, photography is a good tool to accumulate data in a short amount of time. It is also a good support to start morphologic and morphometric analysis using algorithms.

### Automatic morphometric analysis on images

Algorithms for edge detections are particularly widely used in computational biology for characterization, quantification, or feature extractions. They aim at identifying points, lines or outlines in an image. Classical edge detectors proceed to the extraction of images or mesh discontinuities. The result is a 2-bit image where black pixels correspond to the outline and white ones to the background. However, other algorithms are necessary for the objects displaying a simple geometry such as lines, circles, and ellipses, resulting to the common use of two feature extraction techniques: the fitting methods and the generalized Hough transform. The Hough Transform has been introduced by Hough for line detection but it allows also to detect parametric curves in an image, such as circles or ellipses (Ballard 1981; Hough 1962). Each edge point of the image votes for all curves passing through it: the method chooses the curves receiving the most votes. In practice, the space of possible parameters is discretized and the votes are stored in an *accumulator* array: the discretization influences the precision of the method (Duda and Hart 1972; Maitre 1985).

The objective of this present study is to develop a semi-automatic image analysis method using for the first time the Hough transform on jellyfish to approximate the gonads and the umbrella by ellipses. The original step sheet includes: i) implementation of the detection of ellipses by Hough transform in Matlab^®^ (R2017); ii) extraction of the main morphologic and morphometric characteristics of a dataset of jellyfish images; iii) statistical analysis on the morphometric parameters to highlight the discriminant ones between the tetramerous jellyfish and the non-tetramerous ones.

## Materials and methods

### The model, Aurelia

As stated by Dawson and Martin (2001), the moon jellyfish *Aurelia* is among the most widely distributed of all scyphozoans (Mayer 1910; Arai 1997). While the genus was traditionally composed of two species – *A. aurita* (L., 1758) a common inhabitant of nearshore waters circumglobally between about 50° N and 55° S, and *A. limbata* (Brandt, 1835) a polar species – Dawson and Jacobs (2001), analyzing DNA sequences data from nuclear internal transcribed spacer one (ITS-1) and mitochondrial cytochrome oxidase c subunit I (COI), have shown that the genus *Aurelia* included *A. limbata, A. labiata* (L., 1758), and that the species *A. aurita* was a cryptic species, with at least seven distinct clades, several clades warrant recognition as distinct species. If the species inhabiting the Black Sea and the Bosphore, is *A. aurita*, the species inhabiting the waters near Mljet (Adriatica Sea in Croatia) is ‘*Aurelia* sp. 5’. More recently, in a fine paper, Scorrano *et al*. (2017) have analyzed the *Aurelia* species in the Mediterranean Sea. Beside a remarkable morphological analysis of different geographical strains – umbrella shape, diameter and thickness, manubrium shape, gastric pouch shape and size,⃛ –, they distinguished three species: *A. solida* Browne, 1905, *A. coerulea* von Lendenfeld, 1884, and a new species *Aurelia relicta* Scorano, Aglieri, Boero, Dawson and Piraino, 2017. From this paper, the jellyfish inhabiting the Mediterranean coasts of France would be *A. coerulea*. For information, the complete mitochondrial genome of *A. coerulea* completed with a phylogenetic analysis, has been published recently by Seo *et al*. (2020). The situation could be simple if, recently, a study on a jellyfish population in a Tunisian Lagoon, quite close to the Berre Lagoon abiotic parameters, has been identified as *A. solida*, a Lessepssian species, native from the Red Sea (Gueroun *et al*. 2020). Is it the case in the Berre Lagoon? In the present study, in lack of DNA analyses, we rule on the name *Aurelia* spp.

Currently, as the moon jellyfish *Aurelia* is the most frequent jellyfish in demonstration in quite all aquariums of the World – Monaco oceanographic museum, La Rochelle, and Nausicaá in Boulogne (France), Monterey in California (USA), etc. –, and that dispersal events probably influenced the modern zoogeography, we consider, in sight of small morphological even subtle morphological differences, the model, referenced as *Aurelia* spp., in this study, as a representative of the genus *Aurelia*, and that it could be a valid model for the present morphometric analysis.

### The site study

The Berre lagoon on the Mediterranean coast of France, close to Marseille (43°30’N and 5°10’E), is the larger lagoon of the northern coast of the Mediterranean Sea. This large Mediterranean semi-enclosed coastal lagoons (155 km^2^) is connected to the Mediterranean through the long and narrow channel of Caronte. The lagoon is studied since more than three decades, (Kim 1982a,b; Kim 1988; Kim 1998). Since the middle of the XX^e^ century, the size of the lagoon has allowed some heavy industry to develop – oil and chemical companies –, that contribute to pollute water and sediments of the lagoon (Radakovitch *et al*. 2013). Morever, large inputs of freshwaters (more than 6630 · 10^6^ · m^3^ · yr^−1^) and suspended matter released by the EDF hydroelectric power plant since 1966 (Gouze *et al*. 2008), have completely destabilized the ecosystem, that lead away this brackish lagoon eutrophic, exhibiting highly coloured surface waters.

The water quality, and hydrodynamic features is regularly and frequently sampled since 1994 by GIPREB Syndicat Mixte, and some studies work on modeling these parameters (Alekseenko *et al*. 2013; Alekseenko and Roux 2018; Martin *et al*. 2010). The lagoon is 8–9 meters depth, with no stratification in salinity. Periodically, the Berre Lagoon presents strong anoxic crisis as shown in 2018 (GIPREB 2018). For more data we will refer to the thesis of Marchessaux (2019).

In addition, the Berre Lagoon is subject to invasive species, as shown by Marchessaux *et al*. (2017). Among the aquatic invertebrates there are two gelatinous species, *Mnemiopsis leidyi* A. Agassiz, 1860 and *Gonionemus vertens* A. Agassiz, 1862, two invaders that have interfered with the lagoon biocenosis.

### Samples, image acquisition and treatment

#### Sampling procedure

Jellyfish were carefully collected near the shore with a net on March 17^th^, 2006, at Berre-l’Étang harbour, and on May 2^nd^ 2013 at the Jaï beach. All the specimen were fixed in the field in 4% formalin in seawater (i.e. 4 parts formalin [37% w/v] and 96 parts seawater) solution. As the scyphozoans are very fragile and so utmost care must be taken in handling them both before and after fixation to prevent damage. Before image acquisition and treatment, samples were kept suspended in liquid at all times, never handling them out of the water. Specimens preserved in formalin should be stored in sealed containers that are resistant to (or protected from) breakage, in a cool, shaded, well ventilated area.

#### Image acquisition

The jellyfish image acquisition was performed with a Nikon^®^ D800 camera coupled with a Nikkor 35 mm (AF 35 mm *f/*10) lens, and resulted on a 36.3 Mo pixel photography for each specimen (Figure **1**). Considering their *Bauplan*, the *Aurelia* jellyfish specimens were always placed following the flatness of their post-mortem position on a flat, horizontal overhead projector window. A plastic blue background was placed underneath the jellyfish to enhance the contrast and have a better view on the organs – as the organism is transparent.

The main morphological characteristics of interest in this study are the ellipse shape parameters of the umbrella, the number of gastro-gonadic pouches (thenceforward simplified *gonads* throughout the text) and the ellipse shape parameters of each one of them (Figure **1**). A Gaussian filter with standard deviation *σ* = *2*0 was applied in Matlab^®^ (R2017) to each image. They were also resized keeping only 1/20^*2*^ pixels from the original: in the original image, 275 × 275 pixels correspond to 1 cm^2^, in the reduced one, one pixel corresponds to 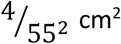. We only test the potential measurement error of total area of the umbrella on a single specimen, taking three pictures after removing it from the glass. The error was lower than 0.05% using ImageJ^®^.

### Gonads and umbrella detection using Hough transform

Since the gonads and the umbrella are not perfect ellipses and the gonads are often not even closed curves, detecting them is non-trivial. The definition of an ellipse *ε* involves five parameters: the center *C* = (*x*_*C*_, *y*_*C*_ ), the orientation *α* ∈ [0, *π*], the semi-major axis *a* and the semi-minor axis *b* (Figure 3). The eccentricity *e* of the ellipse is then given by

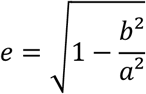

**Figure 3.**
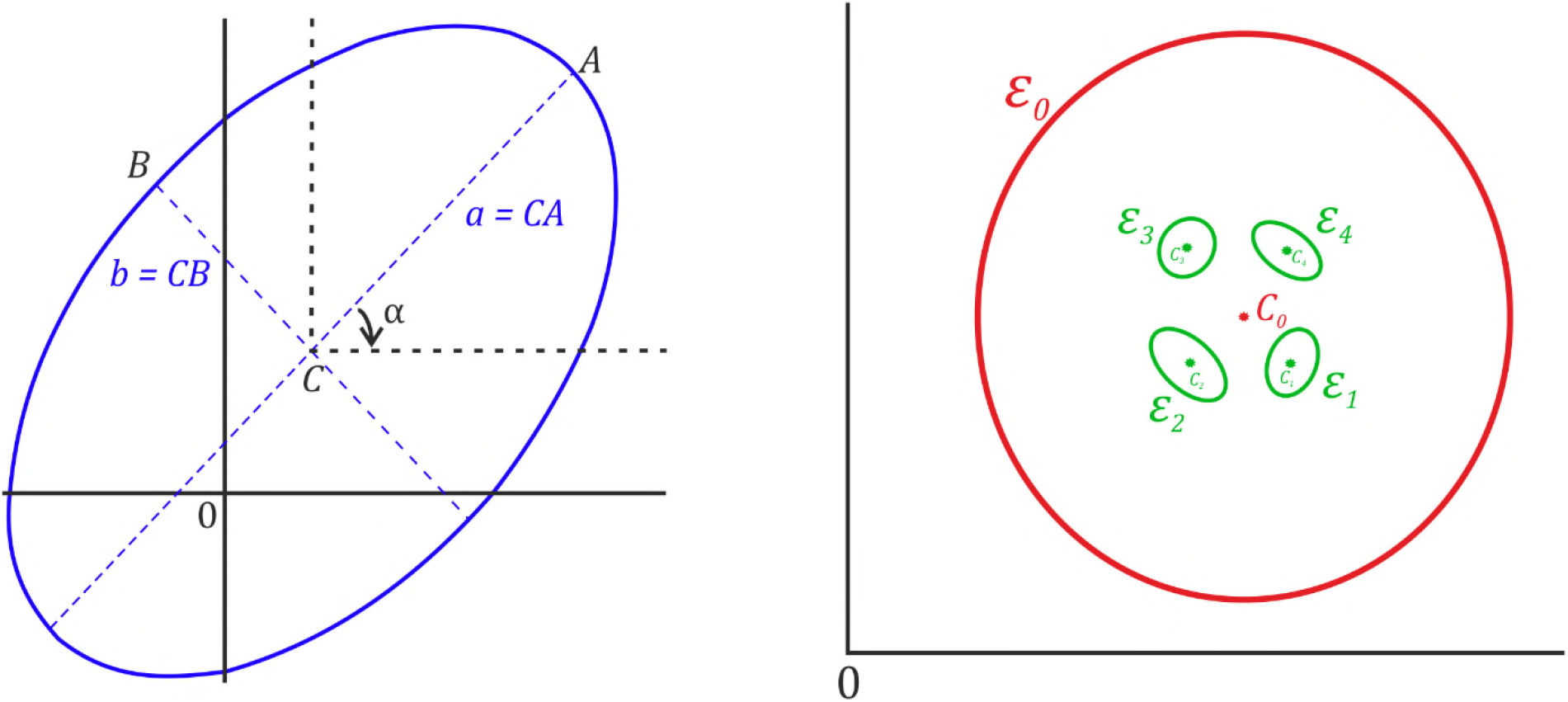
On the left: ellipse *ε* parameters; on the right: ellipses in a tetramerous jellyfish (with N = 4 gonads).

For each jellyfish image, *ε*_0_ denotes the ellipse corresponding to the umbrella, *ε*_1,_ … , *ε*_*N*_ the ellipses corresponding to the *N* gonads (clockwise order) and *C*_*j*_ the center of *ε*_*j*_ (Figure 3).

The implementation of the Hough transform proposed here is based on the parametrization of an ellipse *ε* by its center *C* = (*x*_*C*_, *y*_*C*_ ), its orientation *α*, its semi-minor axis *b*, and its eccentricity *e* ∈ [0,1[. Using these parameters, the equation of *ε* rewrites as

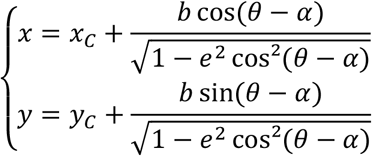

where θ describes [0, 2π].In addition, its major semi-axis *a* is then equal to 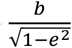

The raw output of the Hough transform is a tensor (array with 5 dimensions) where each cell contains the number of votes obtained for each discrete value of the vector (*x*_*C*_, *y*_*C*_, *α, e, b)* of the 5 parameters. Then it is not possible to visualize this raw output whereas it is for the original Hough transform that detects lines, simply described by 2 parameters (in that case, the raw output is a 2-dimensional array and can then be visualized as a discrete image). The Hough transform finally retains the value of the parameters receiving the most votes. The range of possible parameters is however huge and depends on the size of the image, so that the method can be very slow. To reduce the computational time, the advantage of some *a priori* on the parameters values has been taken. It also allows to avoid some false detections. First the approximate centers of the umbrella and gonads ellipses are set by the user: the detection will focus only on possible centers around these initialized positions, which reduces drastically the possible sets of centers *C* = (*x*_*C*_, *y*_*C*_ ). The size of the neighbourhood is a parameter empirically chosen in function of the object size, leading to choose at least one neighbourhood size for the detection of the umbrella and one other for the detection of the gonads. An *a priori* of the range for the semi-parameters *b* has also been empirically taken; this range can be chosen for each ellipse if needed. Finally, the eccentricity has been bounded by a value *e*_*max*_< 1 to avoid too flat ellipses and false detections. The Hough transform is applied on the set of pixels detected as edge pixels by the Canny edge detector (with the threshold automatically chosen by Matlab^®^ (R2017), in its *Image processing toolbox*). For each image, the mesh size for the orientation *α* is 0.02*π*, the mesh size for the eccentricity *e* is 0.05 and *e* varies between 0 and 0.85: the choice of these mesh sizes is again a compromise between computational time and accuracy of the parameter estimation, and it is again an empirical choice. In addition, the range for the semi-minor axis of the umbrella is [50: 150] (expressed in pixels). For the detection of the gonads, [1:10] has been used as range for the semi-minor axis *b*. On few images, this range was changed for some specific gonads due to their small size: the peaks in accumulator array that should characterize them is not so evident when choosing a comparatively large range for parameter *b*.

The implemented algorithm finally provides each ellipse parameters 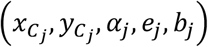 for *j* = 0 …*N*.

### Statistical analysis of some features

Then, several morphologic and morphometric parameters can be computed. Only the significant discriminant morphometric ones highlighting difference between the tetramerous and non-tetramerous jellyfish groups, or giving pertinent morphological clues are exposed in these results. To this aim of comparison, the algorithm computes the following indicators:

- the area of the ellipse *ε*_*j*_

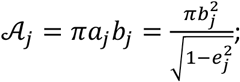
- the ratio (in %) gonad area on area of the umbrella

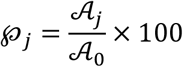

which corresponds to the relative size of the j^th^ gonad with respect to the umbrella one and the associated individual mean area ratio (in %);

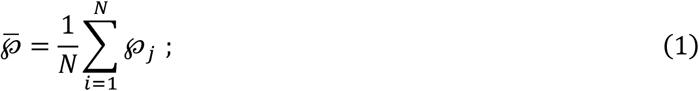
- the ratio

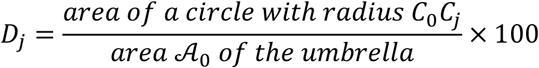

which characterizes the relative distance between the j^th^ gonad and the umbrella center and the associated individual mean ratio (in %);

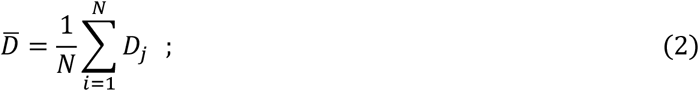
- the individual mean eccentricity

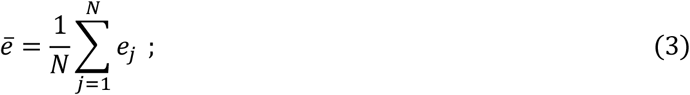
- and the individual variability of the gonad’s eccentricity

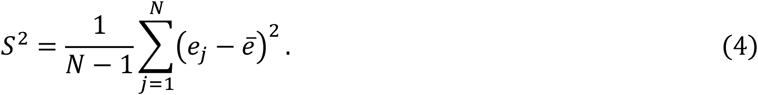

On a given dataset, grouping jellyfish in function of the number of gonads, independent samples of the morphologic and morphometric parameters are thereby obtained. The non-parametric Kruskall-Wallis test can then be performed to test if there is a significant difference according to the number of gonads as regards a chosen parameter. When only two groups are considered, as in our study, this remains to apply Mann & Whitney tests.

## Results

### Dataset

To test the algorithm, the dataset is composed of 19 jellyfish sampled during two proliferations (March 2006 and May 2013) at the same location (Berre Lagoon, south of France). They are divided in 4 groups depending on their number of gonads (Table 2).

**Table 2.** Morphology details of the jellyfish of the data basis.

|  | Tetramerous jellyfish | Non-tetramerous jellyfish |  |  | Total |
| --- | --- | --- | --- | --- | --- |
| Number of gonads | 4 | 3 | 5 | 6 |  |
| Number of jellyfish | 12 | 1 | 5 | 1 | 19 |

The morphologic and morphometric parameters from the jellyfish having 3 and 6 gonads were not included in the statistical analysis because of their very low workforce (*n* = 1 for both of these groups). Let us emphasize that for the statistical study, this leads to compare two independent samples, one of n_1_ = 12 tetramerous jellyfish (*N* = *4* gonads) and one of n_2_ = 5 jellyfish with *N* = *5* gonads. This two samples are very small, and then the statistical part is provided as a preliminary study, and its conclusions have to be relativized.

### Graphic results

Each specimen of the dataset has been treated using a Macbook Air^®^, with Intel i7, 1.7 GHz, and memory 8 Go. Even though our code is not optimized (since written with several nested loops), the computational time is less than two minutes by image. More precisely, since the image sizes are analogous, so that the times for filtering and edge detection are quite the same, around 5-6 seconds. Except for a few pictures, the same ranges for the detection of umbrella and gonads have been used, so that the time for each umbrella detection is around 40–50 seconds and for each gonad around 5–7 seconds.

Figure 4 shows the main morphologic features estimated by the algorithm: the ellipse detected for the umbrella and its center, the ellipses detected for each gonad and their center.

**Figure 4.**
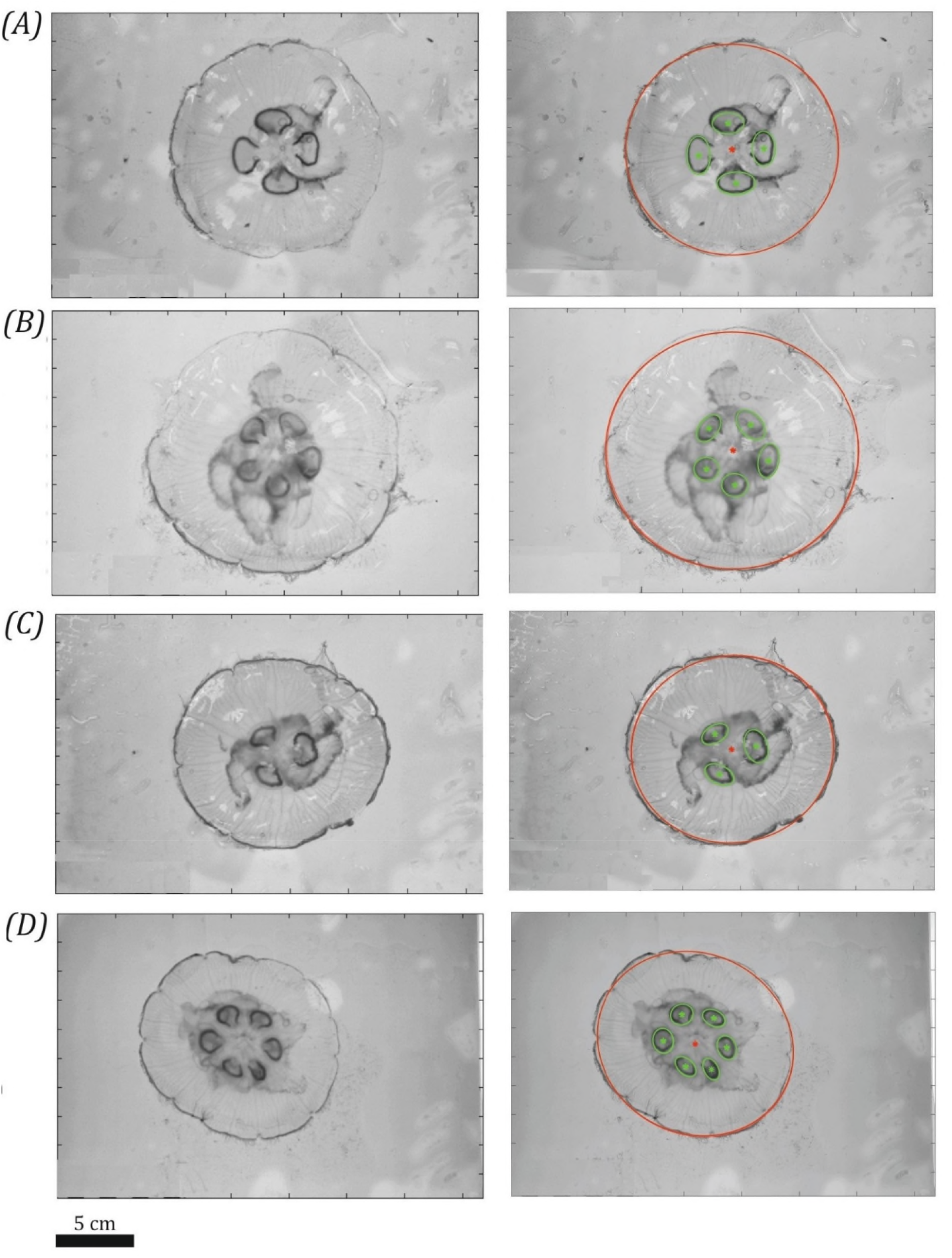
Jellyfish with 4 gonads (A), 5 gonads (B), 3 gonads (C), and 6 gonads (D). Left: grayscale image; right: grayscale image with detected ellipse: red for the umbrella and its center and green for the ellipses detected for each of the *N* gonads.

### Statistical tests

#### Ratio gonad area on area of the umbrella

We tested if for tetramerous and non-tetramerous jellyfish, the relative size of the gonads is the same. To this aim, the individual mean ratio 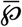 defined by Equation (1) provides a criterion of comparison which does not depend on the organism size. Then the bilateral Mann & Whitney test was used to compare this individual mean ratio for tetramerous jellyfish (*N* = 4) and jellyfish with *N* = 5 gonads. The p-value equals to 0.6461 and so the test leads to conclude that there is no significant difference between the two groups.

#### Distance of gonad center to umbrella center

This part is interested in the location of the gonads, and aims to compare their distance from the umbrella center for tetramerous and non-tetramerous. To take into account the size of the jellyfish and its anisotropy – that is its elliptic shape –, the distance between a gonad and the umbrella center is characterized by the individual mean ratio 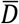 defined by Equation (2): the smaller it is the closer to the umbrella center the gonads are. A unilateral Mann & Whitney test has then been performed in order to test if the individual mean ratio 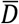 is stochastically greater for jellyfish with *N* = 5 gonads than for the tetramerous jellyfish. The p-value of this test equals to 0.09729, which leads to a weak evidence against the null hypothesis. In other words, this study suggests that for *N* = *5* gonads, with a level significance of 0.1, the gonads of the jellyfish are located further from the umbrella center than in tetramerous jellyfish.

#### The gonad eccentricity

Finally, this part compares the gonad eccentricity between tetramerous and non-tetramerous jellyfish. First, the individual mean eccentricity *ē* defined by Equation (5) has been studied: the bilateral Mann & Whitney test concludes that there is no significant difference between tetramerous jellyfish (*N* = 4) and jellyfish with *N* = *5* gonads as regards the individual mean eccentricity (p-value = 0.2912).

As working on the gonad’s eccentricity, a unilateral Mann & Whitney test has also been performed to compare its individual variability S^2^ for tetramerous jellyfish with the one for jellyfish with *N* = *5* gonads. This test concludes that there is a significant difference between the two groups (p-value = 0.04071): the jellyfish with *N* = *5* gonads exhibit a higher gonad eccentricity variability than the tetramerous ones.

## Discussion

### Implementation of Hough transform in biology

As conspicuous and important component of the ecosystem, medusae have received growing interest over the last decades as numerous proliferations have been reported with increasing frequencies in all seas and oceans. This present study proposes an optimizable tool to quantify and characterize the main *Aurelia* spp. morphometric characteristics through aerial images of *in situ* organisms during a proliferation. Ellipse detection algorithms have been already implemented to characterize circular biological objects such as cells in microbiology microscopic images (Cai *et al*. 2011; Kumagai and Hotta 2012; Lehmussola *et al*. 2005). Here in this paper, it is the first time that the Hough transform has been used to extract morphometric characteristics on jellyfish highlighting new challenge for the implementation such as to detect small ellipses (gonads) in a bigger one (umbrella) and to characterize them as *N* samples of one individual. The algorithm used on cells images has to be optimized in order to be applied to a large sample. In addition, it cannot be directly applied to images in which there is a superposition of ellipses (agglomerated cells), occlusions or two ellipses connected (budding cells). Actually, it forces the biologist to manually set some parameters for a group of images preventing its automation just as here, where the images have to be separated in classes and few parameters fixed to limit the computational time but preventing a full automation. Recently, few authors proposed new solutions such as using gradient accumulation matrix (Denimal *et al*. 2017) or color variation detections by a computer vision system to improve the automation potential (Murillo-Bracamontes *et al*. 2012). Those solutions are currently investigated for an application on aerial jellyfish proliferation captured by drone as shown on Figure 5.

**Figure 5.**
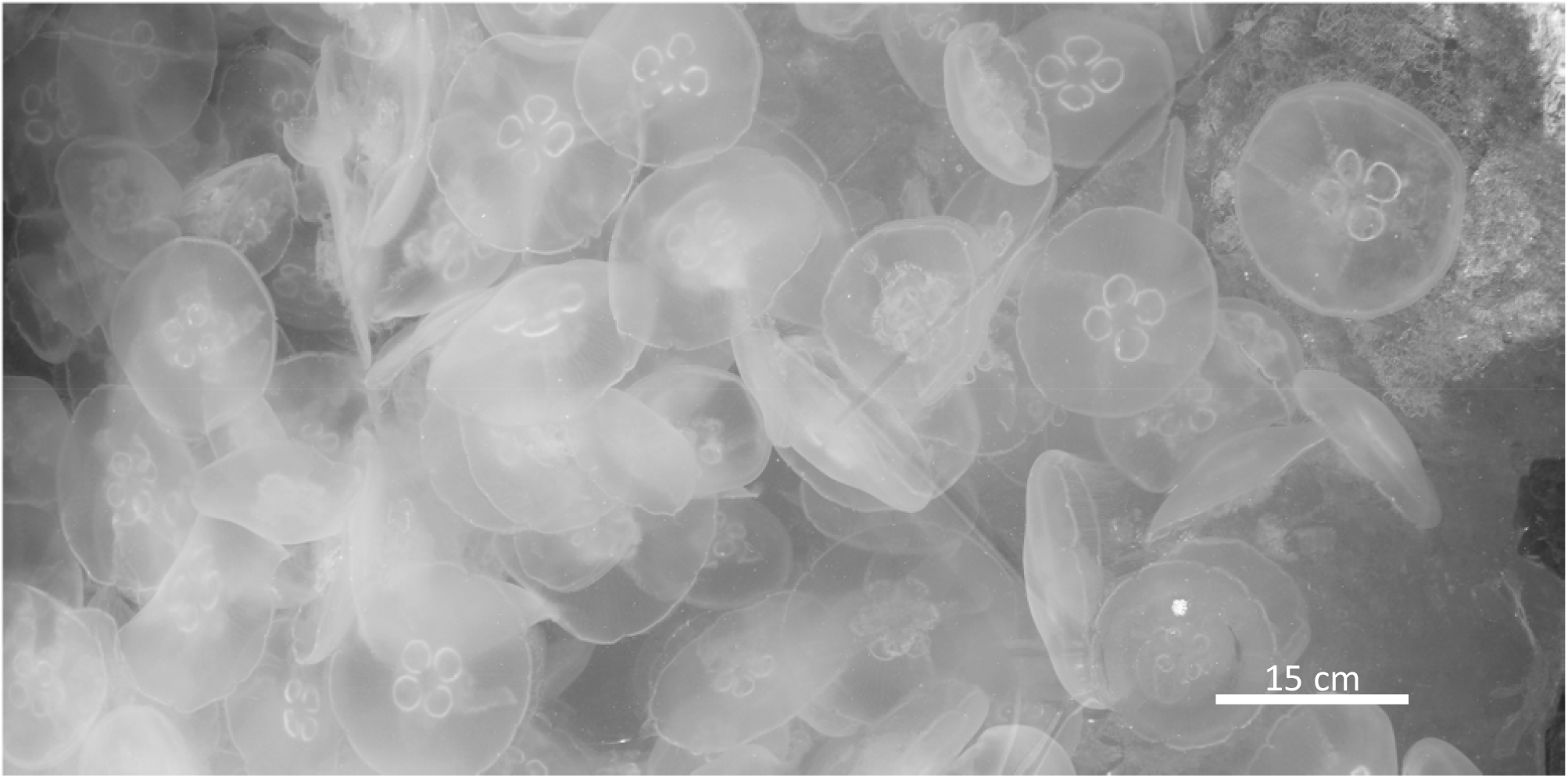
Aerial image of a moon jellyfish *Aurelia* sp. bloom in the the Berre-l’Étang harbour (Southern France) in Sept. 17, 2008 (credit A. Thiéry)

### Discriminant morphometric characteristics

The analysis of the morphometric data has permitted to suggest some remarkable characteristics of the tetramerous jellyfish and the non-tetramerous ones: the ratio 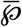 that characterizes the relative size of the gonads, the ratio 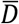 that characterizes the relative distance between the gonads and the umbrella center and the individual variability *S*^*2*^ of the gonad eccentricity (Fig. 6). As already mentioned, this study suggests that the relative size does not depend on the number of gonads, whereas the relative distance and the variability of the eccentricity seem to be higher for jellyfish with 5 gonads. However, its main drawback is that the sample for the tests is very small and so their interpretation have to been relativized: the suggested ecological hypotheses have them to be confirmed using a larger sample. The main force of these analysis is that the results are not population-dependent and organism size-dependent allowing a large application, up to optimizing our code.

**Figure 6.**
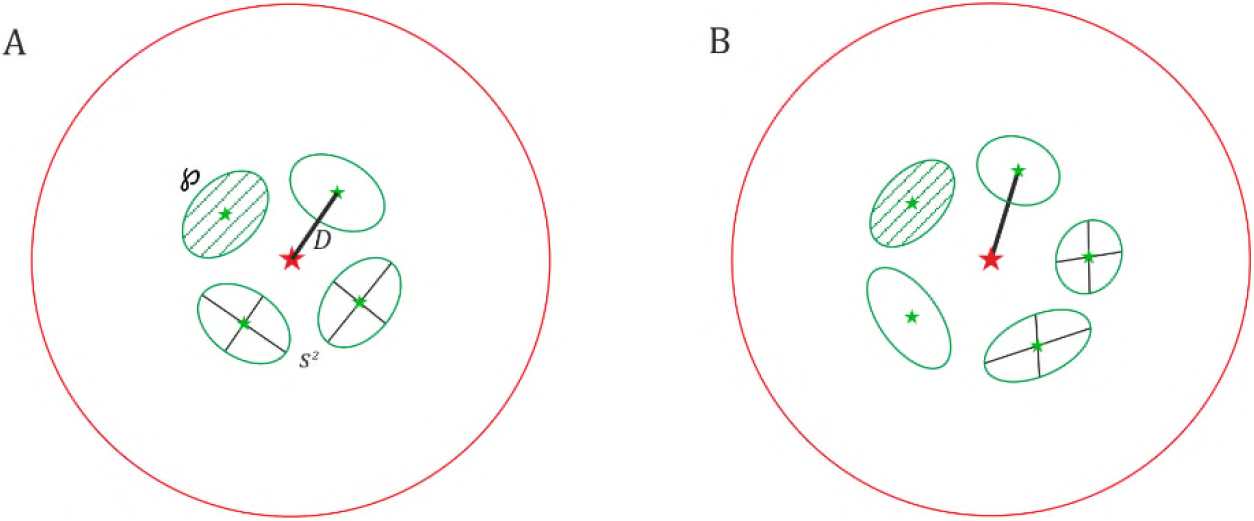
Conceptual scheme of the 3 different morphometric characteristics remarkable between the A: tetramerous jellyfish and B: the non-tetramerous jellyfish. ℘ ratio rendering the relative size of the gonads; S^2^ the individual variability of the gonad’s eccentricity; D ratio rendering the relative distance of the gonad center to the umbrella center.

### Perspective: physiology studies

As the relative size of the gonads seems to be analogous for non-tetramerous and tetramerous jellyfish, the mobility of those dissymmetric jellyfish with more gonads may be affected because the proportion of the muscle in the umbrella is reduced – more surface occupied by the gonads. It may also modify the physiology and the resource allocation: as a reproduction organ, the gonad is a high consumer of energy. Associated with each gonad comes a gastric pouch increasing the potential digestion capacity. More than the number of gonads, the alteration of the tetramerous symmetry also includes the number of *rhopalia* (sensor cells) and sometimes the number of tentacles. Because this dissymmetry appears at a clonal level – for one polyp, normal and dissymmetric *ephyrae* can be produced – the hereditary of these abnormalities is not obvious (Gershwin 1999). In 1999, Gershwin collected a large amount of data through sample analysis and bibliographic reports from 1700s to 1990s. She submitted the hypothesis that the proportion of abnormal development in *Aurelia* spp. can be related to stress, environmental variations, but mainly pollution exposition (Gershwin 1999). This last point was confirmed by the study of Gadreaud *et al*. (2017) in 2017 relating higher proportion of non-tetramerous *ephyrae* born from polyps exposed to emergent xenobiotics, and proposing then this organism as a new model in nanoecotoxicology. Bioassays are currently in progress to test those last results: *Aurelia* sp. polyps are exposed to silver and titanium dioxide nanoparticles which are emergent xenobiotics in marine environment (Gadreaud *et al*. 2017).

In the present times, the ecological monitoring of sensible aquatic ecosystems becomes necessary. As several studies dealing with the physico-chemical and biological parameters monitoring – colour, eutrophication, suspended particulate matter, water turbidity, chlorophyll-a, etc. – by satellite (SWOS Satellite-based Wetland Observation Service), aerial hyperspectral imagery – Projects HYPERBERRE and DCS4COP (Faure *et al*. 2018; Faure *et al*. 2020) –, we propose to complete this basket of solutions, with a biomonitoring of teratological jellyfish in the lagoon. For it we are working with the Society Cambulle^®^(*https://www.societe.com/etablissement/cambulle-48528274300015.html*) society specialized in aerial drone photography, that is capable to capture and store sharp images of jellyfish (up to 500 µm precision) *in situ*, as in Fig. 5. These images will be treated automatically with the present algorithm, even in the case of noisy images as studied by Cohen and Toussaint (1977).

## Supplementary material

### Code details

**Algorithm 1** Detection of the umbrella and the gonads

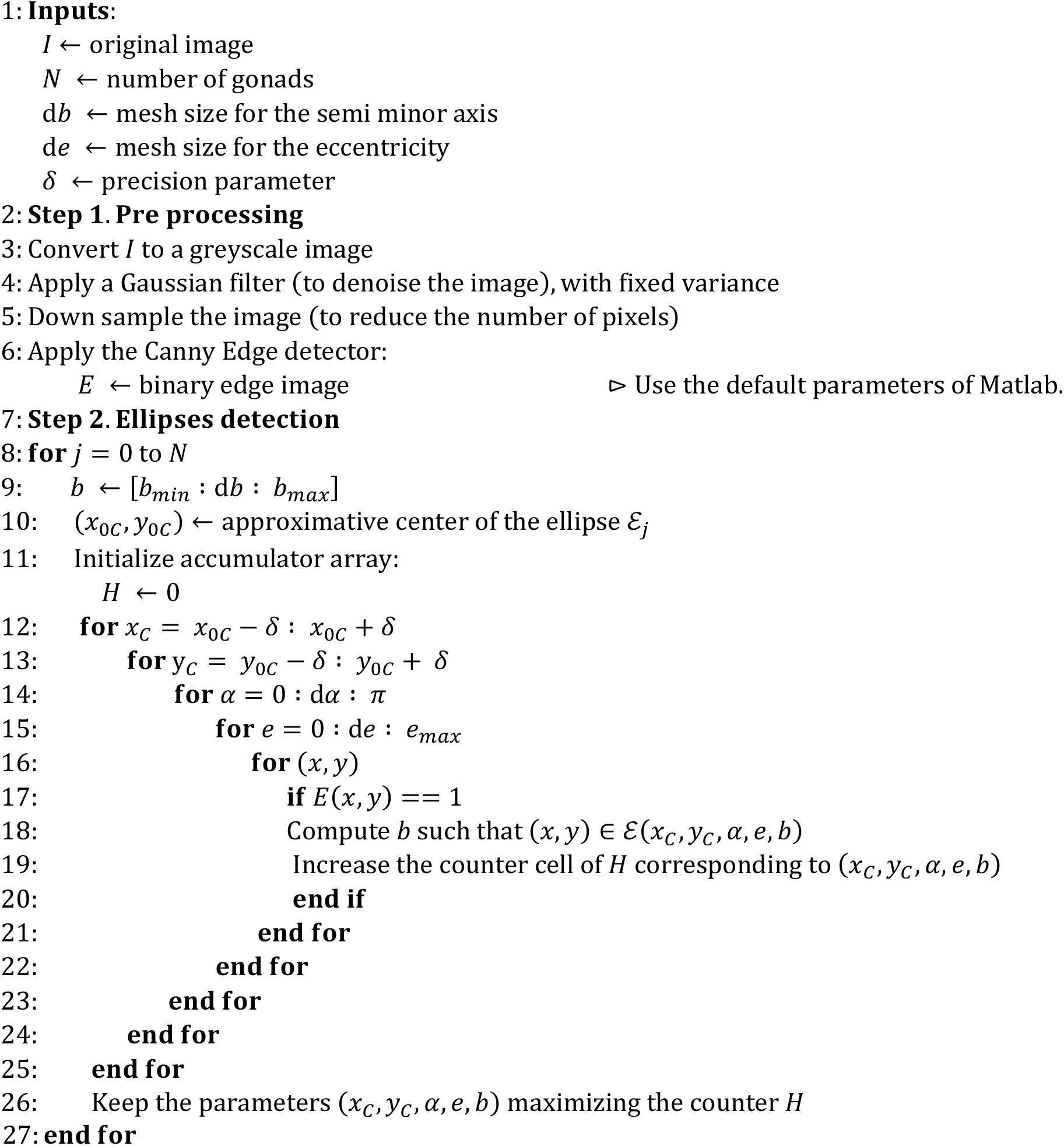

Statistical script in R for Mann & Whitney tests

\# Pmean4, Pmean5 are respectively the sample of the mean individual ratio characterizing the # relative size of the gonads for tetramerous jellyfish and for jellyfish with 5 gonads

\# Bilateral Mann & Whitney test comparing theses two samples wilcox.test(Pmean4, Pmean5,alternative=”two.sided”)

\# Dmean4, Dmean5 are respectively the sample of the mean individual ratio characterizing the # relative distance between the gonads and the umbrella center for tetramerous jellyfish and # for jellyfish with 5 gonads

\# Unilateral Mann & Whitney test comparing theses two samples wilcox.test(Dmean4, Dmean5,alternative=”less”)

\# Emean4, Emean5 are respectively the sample of the mean individual eccentricity of the # gonads for tetramerous jellyfish and for jellyfish with 5 gonads

\# Unilateral Mann & Whitney test comparing theses two samples wilcox.test(Emean4, Emean5, alternative=”two.sided”)

\# S4, S5 are respectively the sample of the individual variability of the eccentricity of the gonads # for tetramerous jellyfish and for jellyfish with 5 gonads

\# Unilateral Mann & Whitney test comparing theses two samples wilcox.test(S4, S5, alternative=”less”)

## Acknowledgements

The authors thank Vincent Bonhomme, Julien Claude, and an anonymous reviewer for comments that improved the article.

This work was supported by a grant of the French Ministry of Higher Education and Scientific Research, and has received funding by the French Anses (Agence nationale de sécurité sanitaire de l’alimentation, de l’environnement et du travail – DecoNano programme 2015 *‘Systèmes bioinspirés pour la décontamination des nanoparticules’*). This preprint has been peer-reviewed and recommended by Peer Community In Ecology (https://doi.org/10.24072/pci.ecology.100055). Version 3 of this preprint has been peer-reviewed and recommended by Peer Community In Ecology (https://doi.org/10.24072/pci.ecology.100066).

## Appendix

### Main program – Matlab code

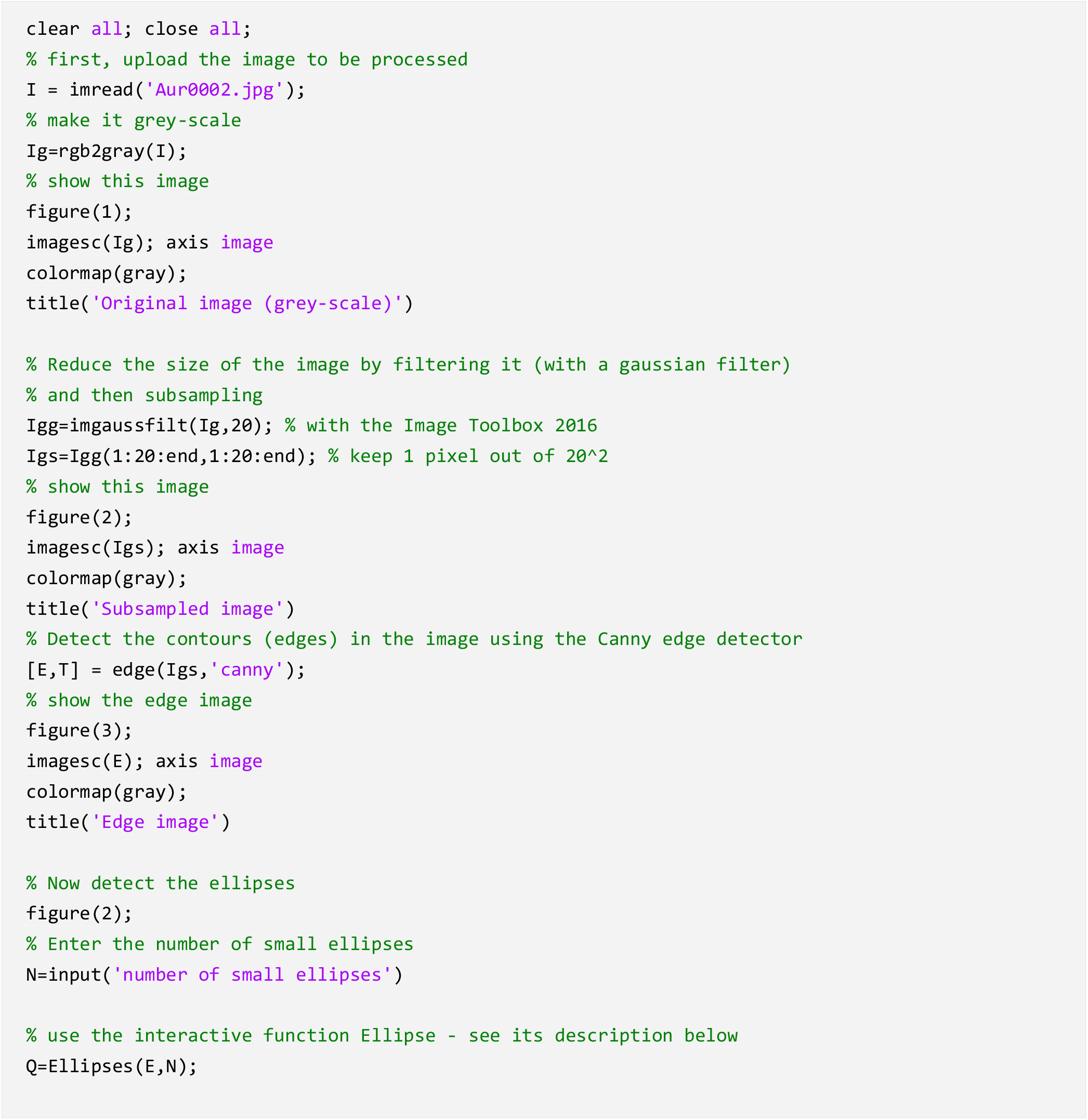
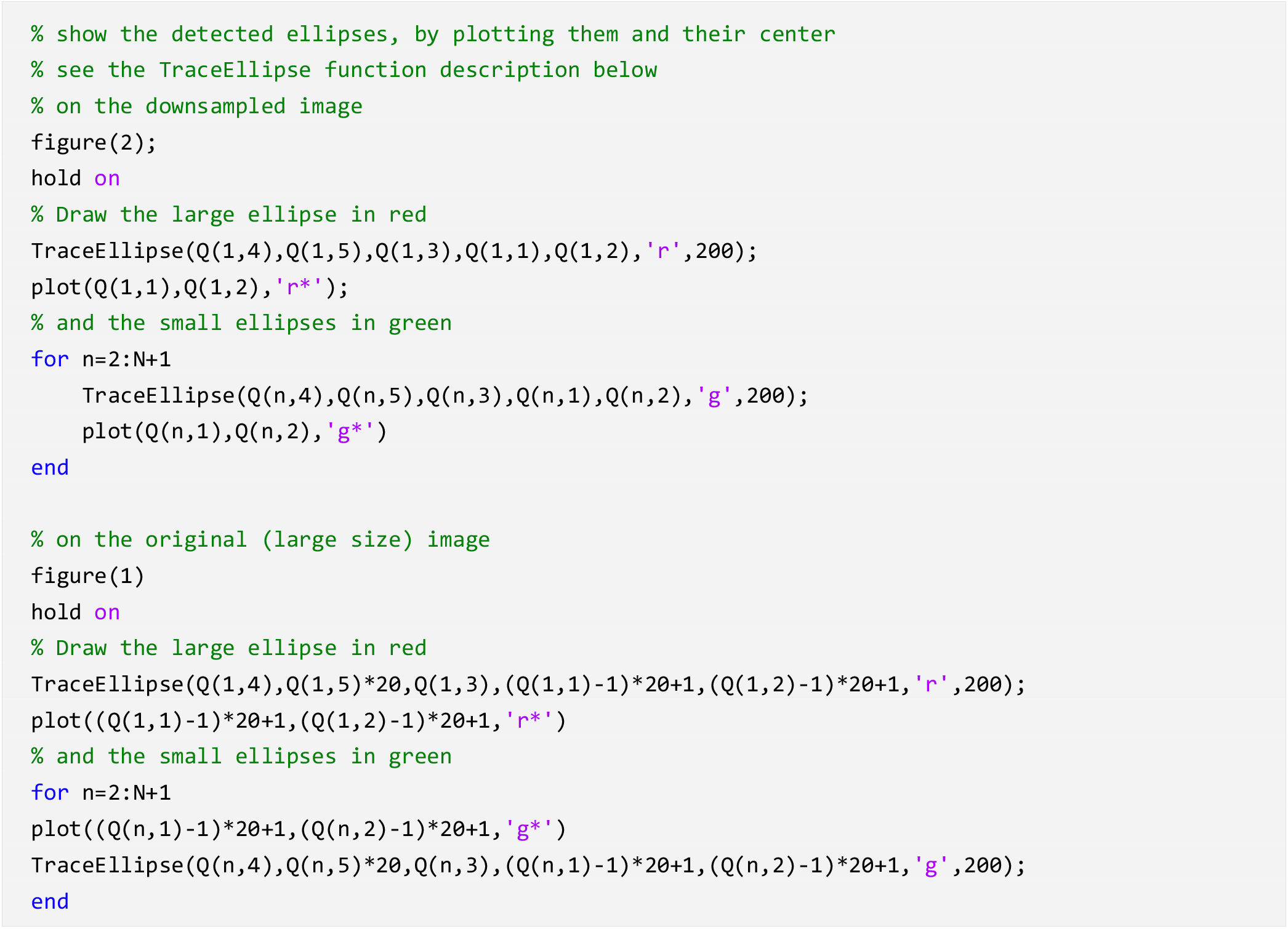

#### Statistical elements of the ellipses

We compute here several quantities on the ellipses in order to have a statistical analysis of these elements

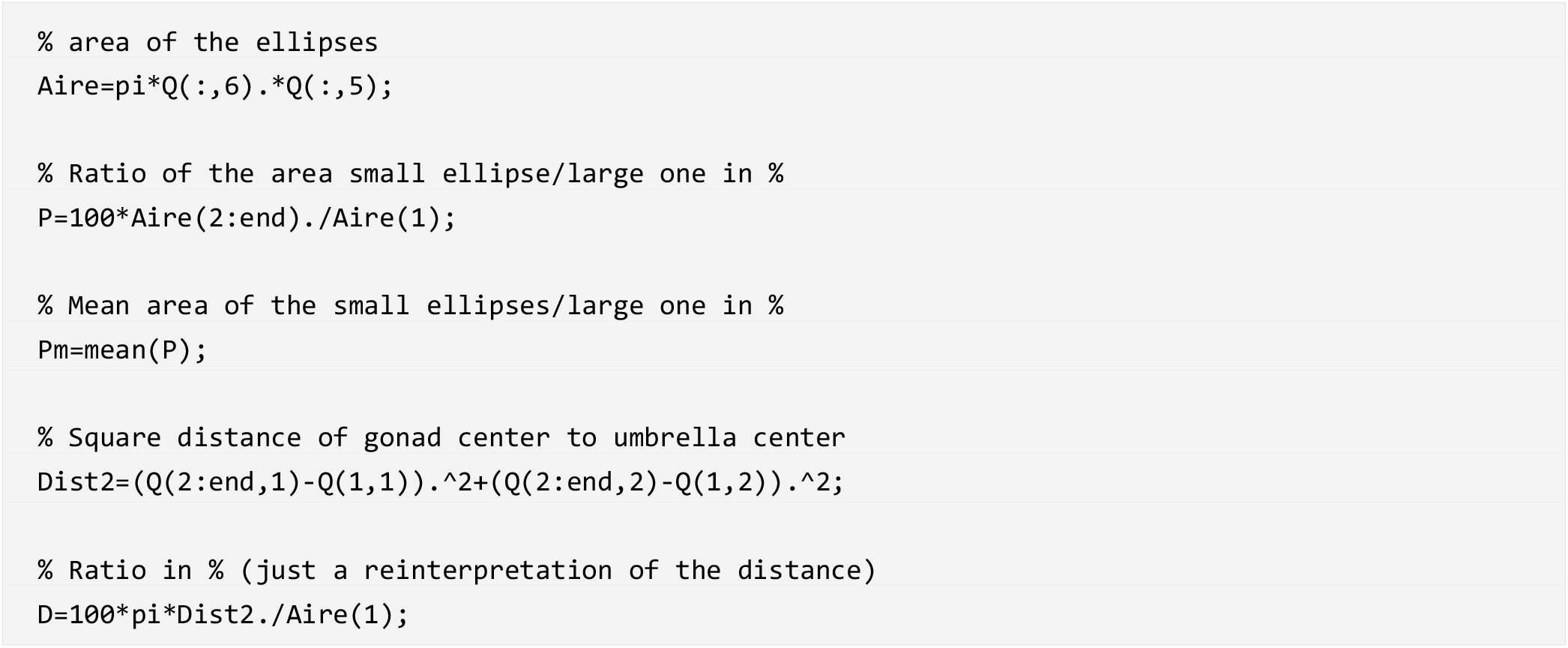
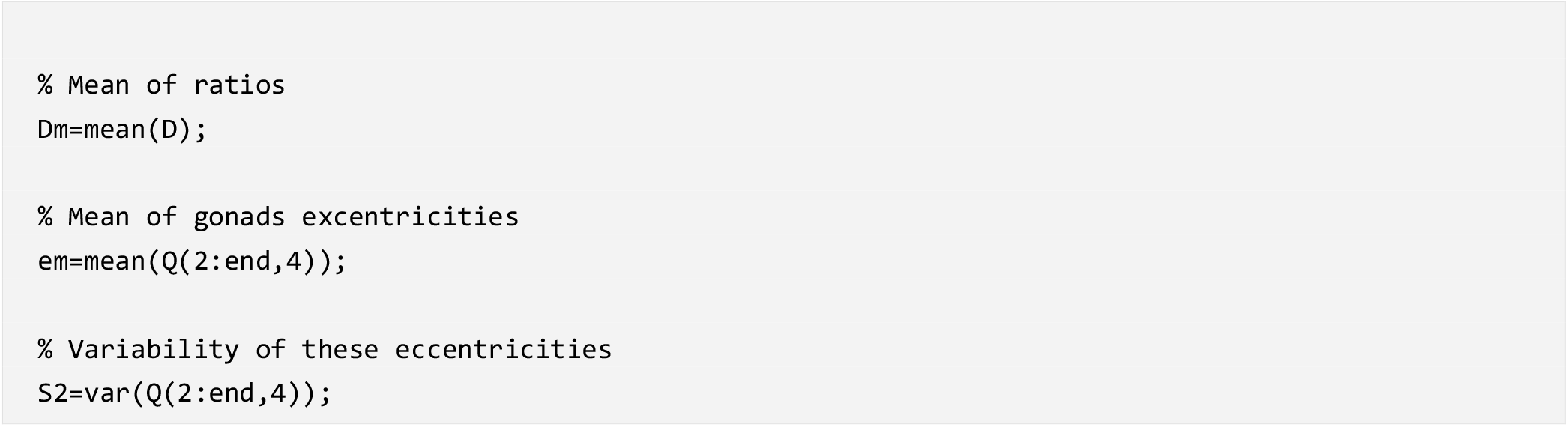

#### Ellipse function

*N* is the number of small ellipses the output Q is an array made of N+1 rows, and row i contains the features of the i-th ellipse: coordinates of the center, orientation, eccentricity, semi-minor axis, semi-major axis

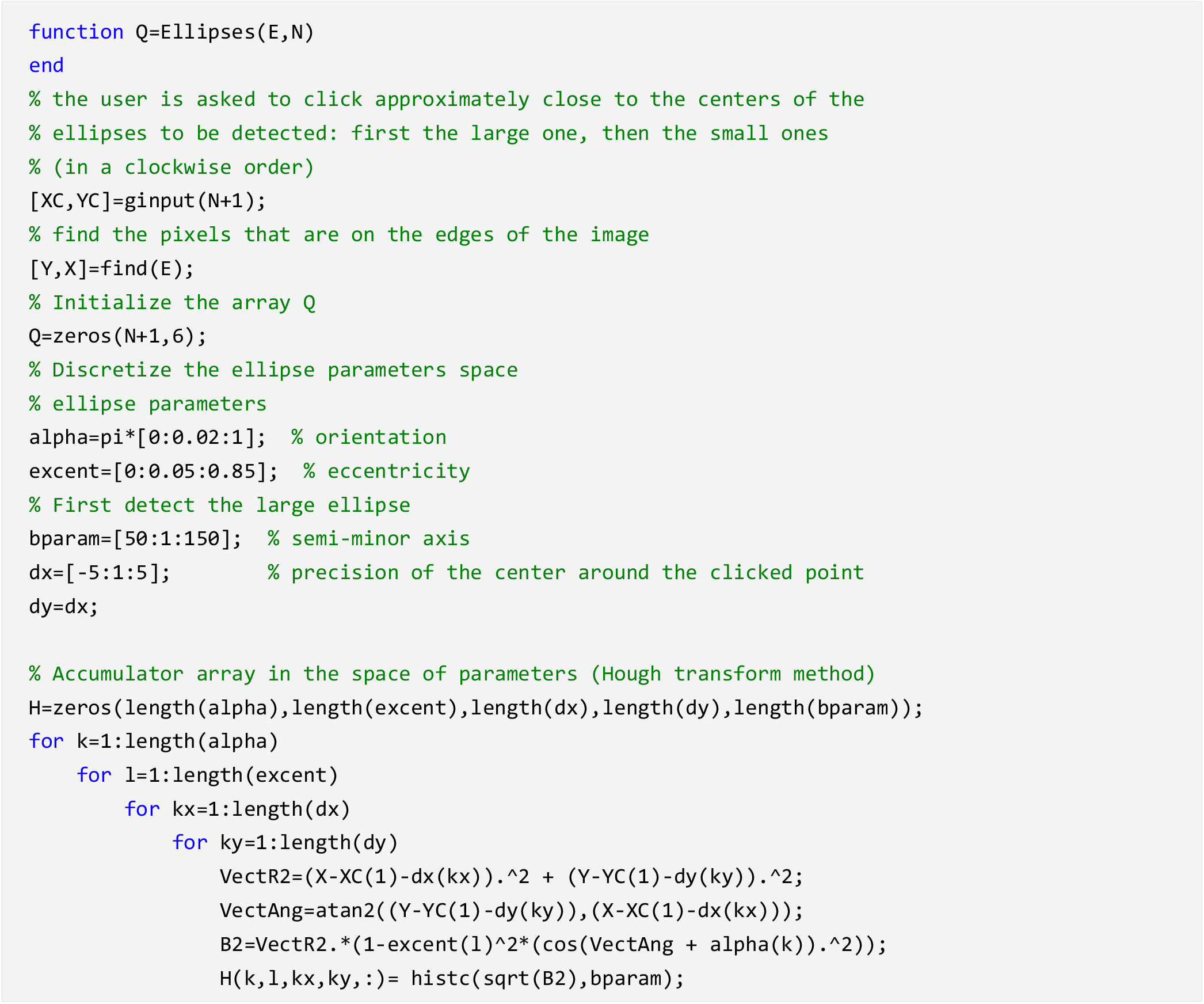
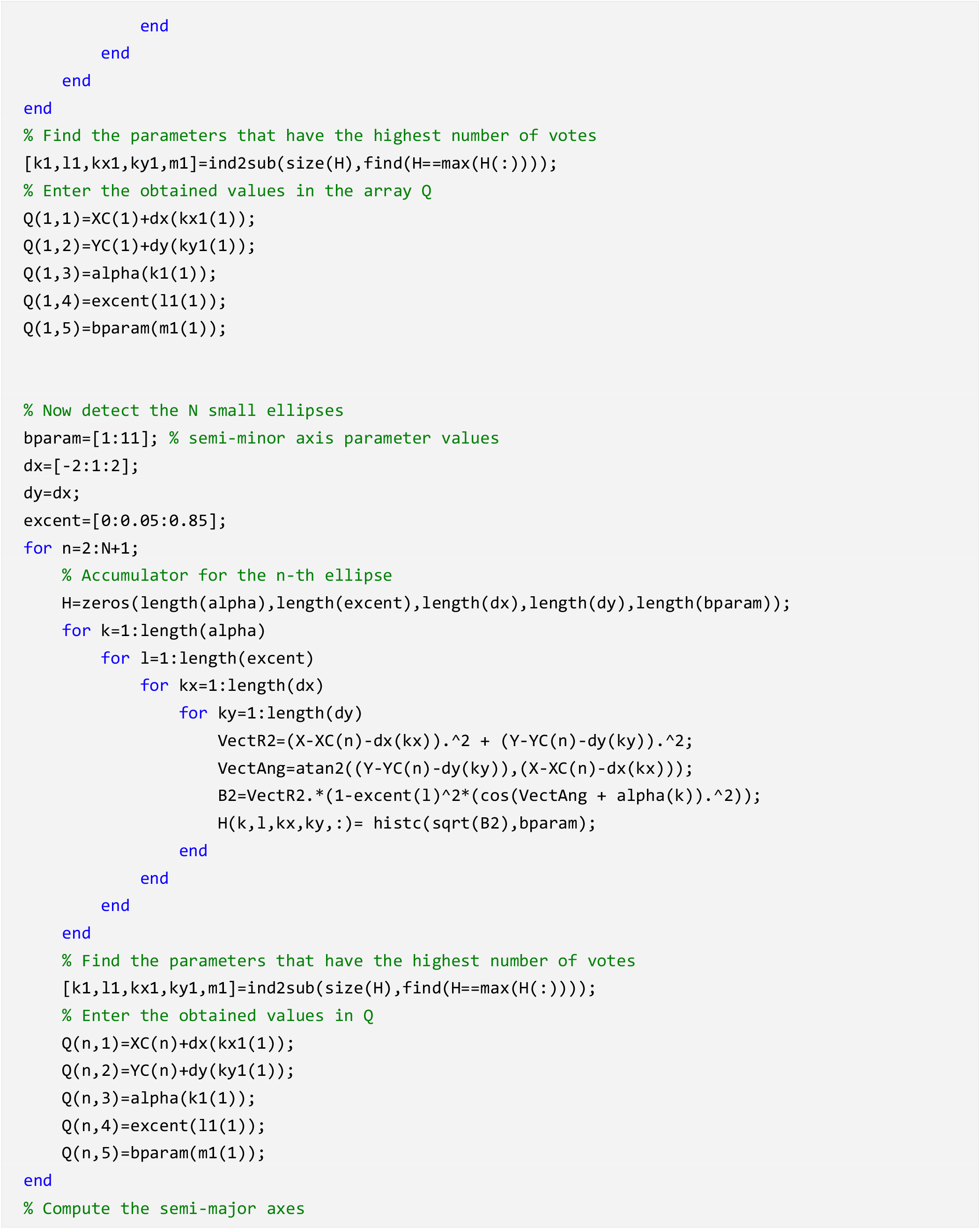
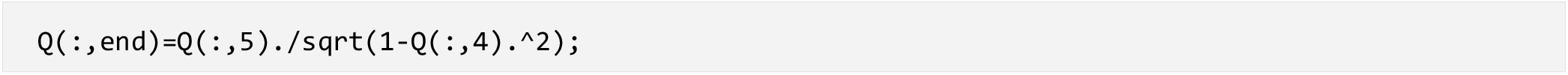

#### TraceEllipse function

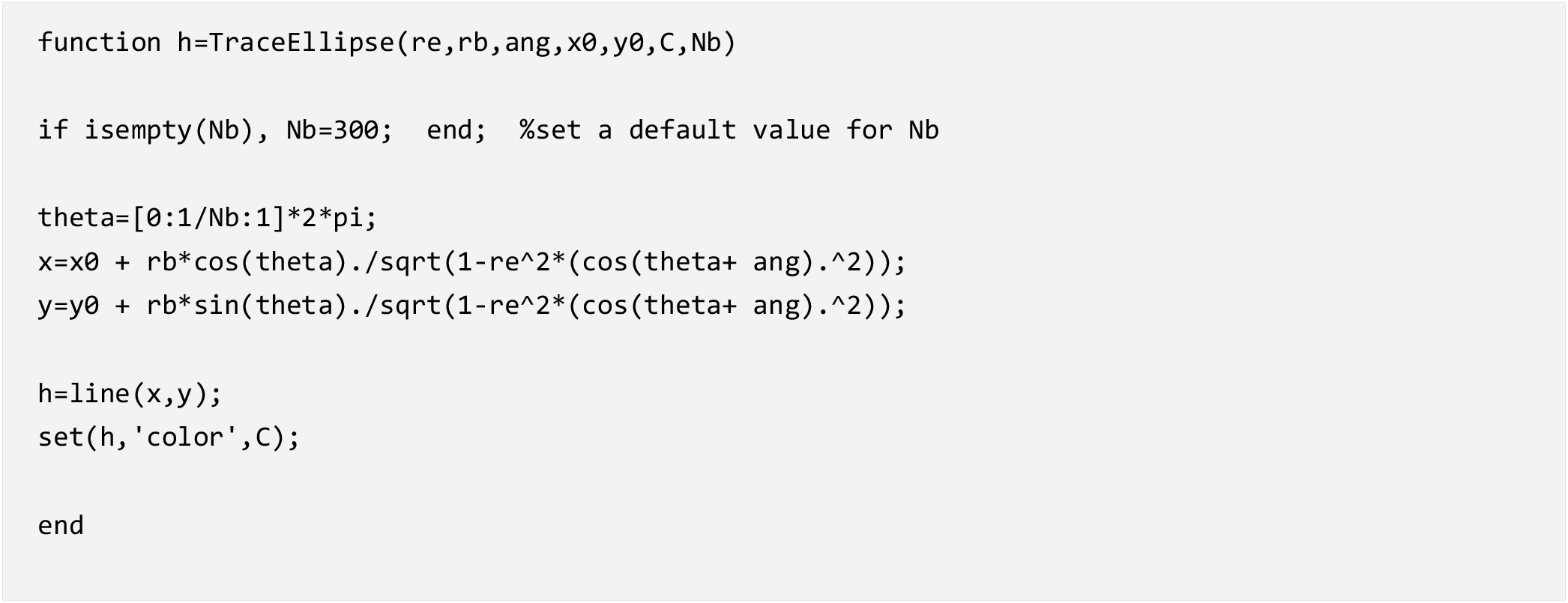

#### Dataset

##### Ratio 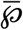

Ratio $\bar{\rho}$
| $N = 4$ | $N = 5$ |
| --- | --- |
| 1,19070475 | 1,10555175 |
| 1,3371931 | 1,64398263 |
| 1,37615931 | 1,30249929 |
| 1,35883684 | 1,33182187 |
| 1,81171429 | 1,49865162 |
| 1,36825379 |  |
| 1,34553927 |  |
| 1,31292481 |  |
| 1,63672109 |  |
| 1,94626611 |  |
| 1,01781433 |  |
| 1,33366571 |  |

##### Ratio 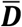

Ratio $\bar{D}$
| $N = 4$ | $N = 5$ |
| --- | --- |
| 7,85372029 | 7,56035905 |
| 8,23246861 | 9,22302216 |
| 8,08573724 | 8,82905582 |
| 9,20822754 | 8,0996472 |
| 8,92085025 | 9,82763814 |
| 7,13892147 |  |
| 7,09617206 |  |
| 7,60209822 |  |
| 6,91098974 |  |
| 9,64754433 |  |
| 6,38226368 |  |
| 8,32379734 |  |

##### Eccentricity

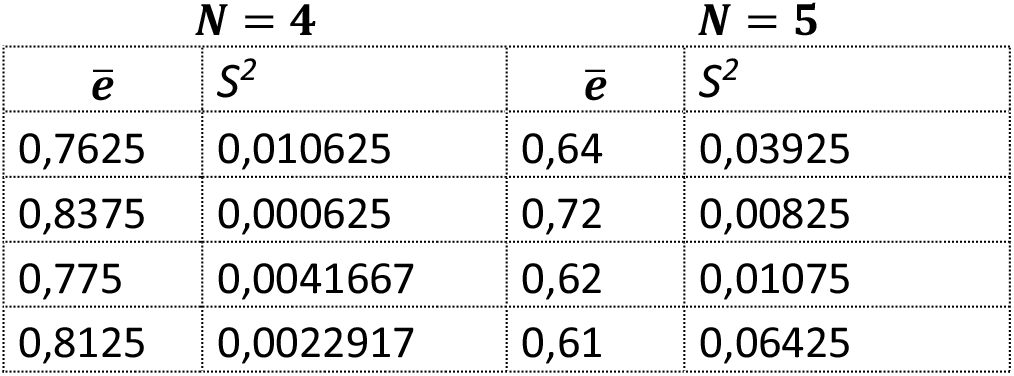
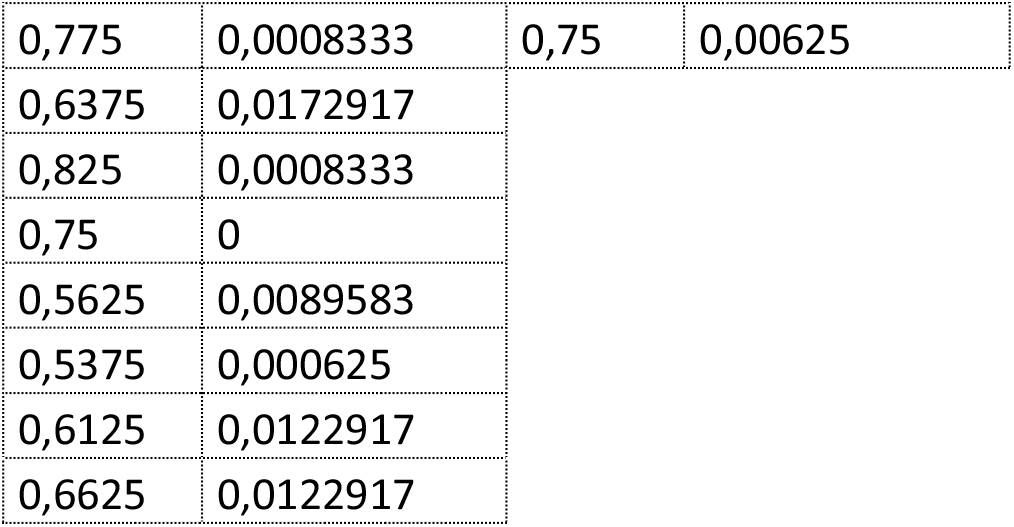

| $N = 4$ | | $N = 5$ | |
| --- | --- | --- | --- |
| $\bar{e}$ | $S^2$ | $\bar{e}$ | $S^2$ |
| 0,7625 | 0,010625 | 0,64 | 0,03925 |
| 0,8375 | 0,000625 | 0,72 | 0,00825 |
| 0,775 | 0,0041667 | 0,62 | 0,01075 |
| 0,8125 | 0,0022917 | 0,61 | 0,06425 |
| 0,775 | 0,0008333 | 0,75 | 0,00625 |
| 0,6375 | 0,0172917 |  |  |
| 0,825 | 0,0008333 |  |  |
| 0,75 | 0 |  |  |
| 0,5625 | 0,0089583 |  |  |
| 0,5375 | 0,000625 |  |  |
| 0,6125 | 0,0122917 |  |  |
| 0,6625 | 0,0122917 |  |  |

